# CulebrONT: a streamlined long reads multi-assembler pipeline for prokaryotic and eukaryotic genomes

**DOI:** 10.1101/2021.07.19.452922

**Authors:** Julie Orjuela, Aurore Comte, Sébastien Ravel, Florian Charriat, Tram Vi, François Sabot, Sébastien Cunnac

**Author notes:** Equal contributor (Julie Orjuela, Aurore Comte, Sébastien Ravel).

## Abstract

Using long reads provides higher contiguity and better genome assemblies. However, producing such high quality sequences from raw reads requires to chain a growing set of tools, and determining the best workflow is a complex task.

To tackle this challenge, we developed CulebrONT, an open-source, scalable, modular and traceable Snakemake pipeline for assembling long reads data. CulebrONT enables to perform tests on multiple samples and multiple long reads assemblers in parallel, and can optionally perform, downstream circularization and polishing. It further provides a range of assembly quality metrics summarized in a final user-friendly report.

CulebrONT alleviates the difficulties of assembly pipelines development, and allow users to identify the best assembly options.

## Introduction

Third-generation sequencing technologies, namely Pacific Biosciences (PB) and Oxford Nanopore Technologies (ONT), provide reads up to 25kb in length, and even hundreds of thousands of bases for ONT. They can extend over repeats or structural variants, and thus result in higher contiguity and accuracy of genome assembly. Due to its low price, speed, portability and easy-access system, ONT is increasingly used worldwide, generally by laboratories sequencing their favorite organism, but with limited expertise in assembly methods. Nonetheless, assembly is not trivial: eukaryotic genomes assembly is a highly complex task (large genome sizes, high rates of repeated sequences, high heterozygosity levels and even polyploidy), and while prokaryotic genomes may appear less challenging, specific features such as circular DNA molecules, must be taken into consideration to achieve high quality assembly.

Numerous long-read assembly and post-assembly tools are available, relying on a large variety of approaches and algorithms, and many of them are frequently updated. However, even with this plethora of tools, there is no single silver bullet to genome assembly for all taxonomic groups. The systematic assessment of these assembly tools is needed to properly exploit data (Murigneux, Rai, et al., 2020). But performing benchmarks to find the best combination of tools for a given dataset and application is a highly complex task (see for example Wick and Holt, 2021 and Chen et al., 2020), even if the computer code used for such studies is increasingly being made available (e.g. Wick, Judd, et al., 2021 or Latorre-Pérez et al., 2020). This endeavor would presumably benefit from streamlined generalist data processing workflows that are accessible, scalable, traceable and reproducible.

A few solutions, such as Katuali (*https://nanoporetech.github.io/katuali/* 2018) or CCBGpipe (Liao et al., 2019), were previously developed to tackle these issues, but they are dedicated to either eukaryotic or prokaryotic genomes only, or provide a restricted choice of assemblers. Some of these are also difficult to parallelize in a HPC environment, or to update with the latest versions of software components. In addition, for the subsequent evaluation of the assemblies, it is critical to organize and aggregate the numerous Quality Control (QC) metrics generated by various tools, in order to facilitate comparisons. As an example, microPIPE (Murigneux, Roberts, et al., 2021), a recently released Nextflow-based bacterial ONT reads assembly pipeline, fills some of these gaps but does not incorporate QC analysis apart from QUAST. To address these issues, we developed CulebrONT, a pipeline allowing users to easily assemble, circularize (if needed) and polish assemblies on multiple datasets with multiple alternative tools, while reporting various QC metrics for each assembly.

### Implementation

CulebrONT assembles, circularises, polishes and corrects genome sequences from raw read sequences in fastq format, and provides QC metrics. To provide more flexibility to the user, individual tools are optional and include the most popular ones. Thus, CulebrONT can be used either to prototype and explore assemblies on new organisms or in assembly production. While originally developed primarily for working on ONT data, CulebrONT can also be used on PB data with generalist tools.

Six recent community-validated assemblers are currently included (Canu (Koren et al., 2017), Flye (Kolmogorov et al., 2019), Raven (Vaser and Šikić, 2020), MiniAsm (Li, 2016) (coupled with minipolish (Wick and Holt, 2020) for an initial polishing), Shasta (Shafin et al., 2020) and smartDeNovo (Liu et al., 2020)). Several tools for polishing (Racon (Vaser, Sović, et al., 2017), Pilon (Walker et al., 2014), Medaka (*https://github.com/nanoporetech/medaka* 2018) and Nanopolish (Loman et al., 2015)) were also included. If requested, Circlator (Hunt et al., 2015) can automatically circularise the primary output of assemblers that cannot handle circular molecules (see Additional file 1 section 2.2 for more details). Each processing step in CulebrONT, from raw to final assembly, can be optionally assessed by various quality control tools (BUSCO (Simão et al., 2015), QUAST (Gurevich et al., 2013), BlobTools (Laetsch and Blaxter, 2017), KAT (Mapleson et al., 2017), Assemblytics (Nattestad and Schatz, 2016), Samtools Flagstats (Li et al., 2009), Mauve (Darling et al., 2004)) and Merqury (Rhie et al., 2020).

CulebrONT uses Snakemake (Köster and Rahmann, 2012) principles and functionalities, enabling readability of the code, local and HPC scalability, reentry, reproducibility and modularity. The pipeline code is written following the Snakemake syntax, and include a dedicated Python module which should facilitate readability for developers. The Snakemake system implements scalability at the level of jobs that are scheduled independently to run, in parallel or sequentially depending on the available resources and the computing infrastructure. In addition, for rules that run programs with internal parallel processing capabilities, it is possible to configure the number of cores allocated to the corresponding individual jobs in CulebrONT. Snakemake handles job failures and a restarted instance of the pipeline will compute only missing output.

CulebrONT relies on conda (*Anaconda Software Distribution* 2021) and singularity (Kurtzer et al., 2017) that simplify installation of specific versions of the software, secure environments and greatly improve reproducibility. CulebrONT is available as a Python Package in PyPi to ease its installation https://pypi.org/project/culebrONT/. In addition, the CulebrONT API adapts the installation to local or distributed/HPC environment. In terms of modularity, relevant tools or processing steps can be selected or omitted in the user setup file *config*.*yaml*, and CulebrONT builds a dedicated instance of the workflow. Data can come from a single sample as well as from multiple ones (one fastq file per sample in the data folder). Tools paths are imported from *tools_path*.*yaml*, and cluster resources are managed using the *cluster_config*.*yaml* file (Figure 1.1). The dedicated CulebrONT class checks configuration files, controls if data and software environments exist and ensures the global coherence of the requested steps. This python class imports snakefiles to build a specific instance of the workflow (Figure 1.2). Upon execution (on HPC or a single machine, Figure 1.3), in addition to the expected final output, individual rules will generate log files (Figure 1.4).

**Figure 1.**
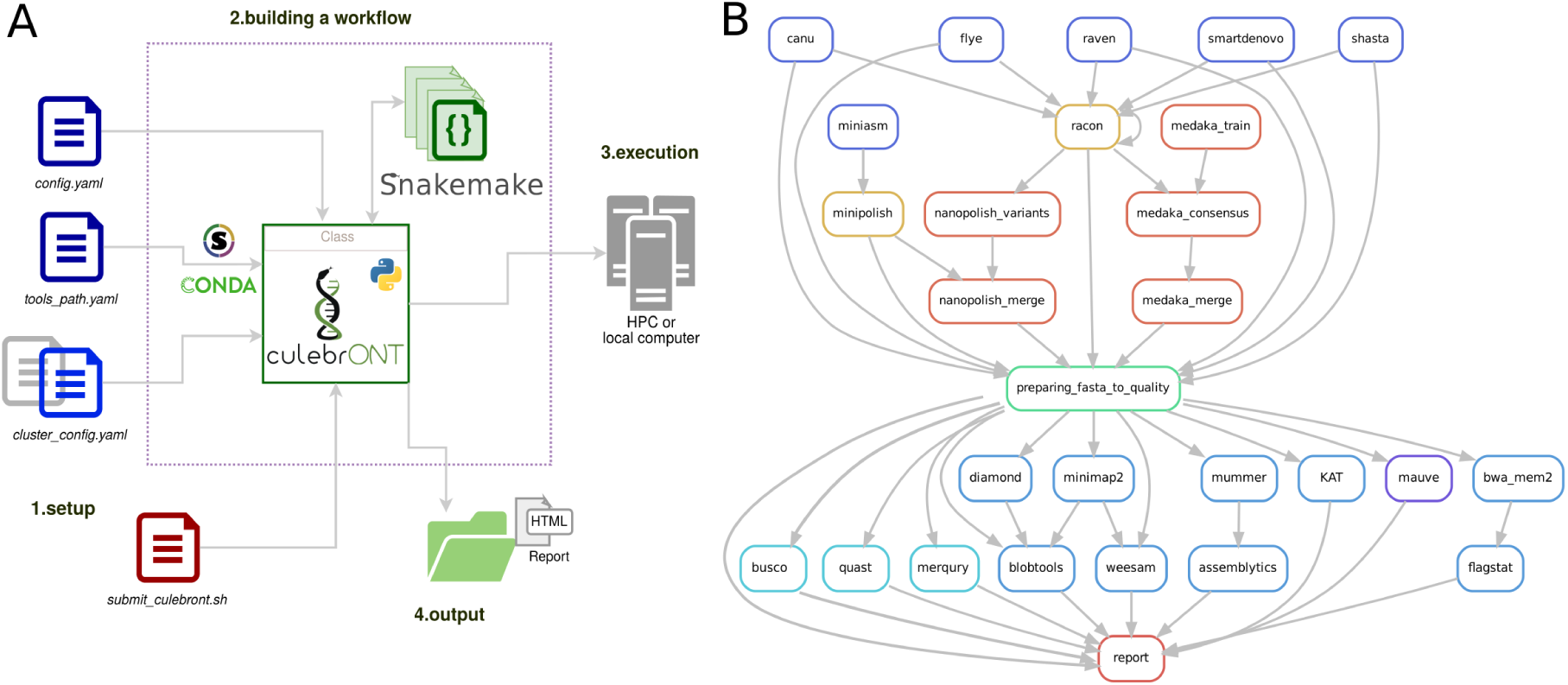
A graphical overview of the CulebrONT pipeline features. **A**. A conceptual diagram of the various tasks involved in setting up a custom CulebrONT workflow: setting up workflow steps and parameters, executing and viewing results of CulebrONT. (1) workflow steps and parameters are provided by the user in the *config*.*yaml, tools_path*.*yaml* and *cluster_config*.*yaml* files. (2) CulebrONT python class reads, transforms and chains rules to build the corresponding pipeline. (3) Rule execution can be performed sequentially or in parallel, on a single machine or a HPC infrastructure, with concurrent jobs and multi threaded tasks. (4) The results of individual steps are aggregated in a convenient HTML output report. **B**. Simplified diagrams showing the components of an example CulebrONT workflow for assembling a linear genome. In the upper part, the tools used for assembly (blue shape outline) and polishing/correction (yellow and red shape outline) are connected with arrows following the directional flow of the pipeline. In the central part, the various versions of the assembled genomes are gathered (*preparing_fasta_to_quality*) to serve as input for subsequent quality evaluation tools in the lower part. Ultimately, the output is aggregated in a final report.

**Figure 2.**
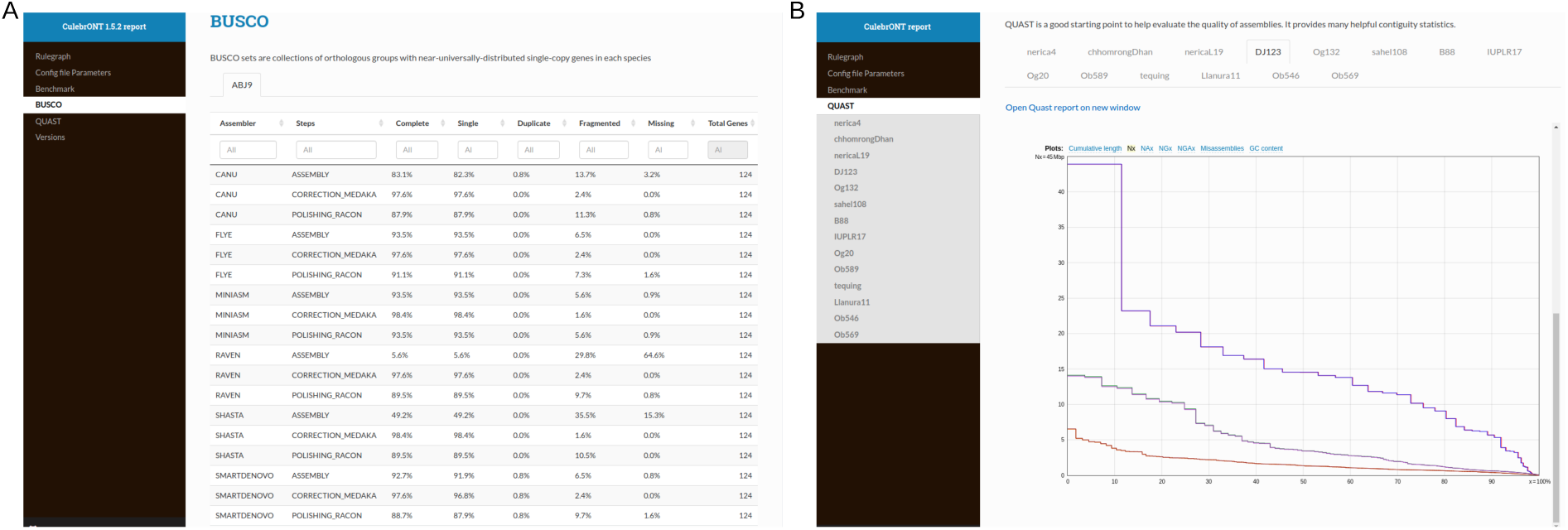
Example of CulebrONT HTML reports. A. Busco statistics for *Acinetobacter sp*. dataset B. Quast for the rice dataset.

## Results

CulebrONT was developed in order to facilitate comparisons between primary assemblies, but also to check the effect of polishing on sequence accuracy. CulebrONT compiles information from these steps, but also essential quality information calculated at each activated step. An individual output directory with a specific subdirectories topology (see Additional file 1 to details) is generated for each samples. It notably contains a sub-directory corresponding to each activated step on the *config*.*yaml* file (assemblers, circularisation, polishing, correction, fixstart, and QC, for instance). In addition, a log directory includes execution information for all steps. CulebrONT generates an global HTML report, found in the *FINAL REPORT* folder. In this report, summarized statistics and relevant information can be found, such as the configuration parameters used or the tools versions and the computational time for each steps (Figure 2). These information can be useful for users that want to benchmark and identify suitable workflows for their data.

In order to provide an illustration of how CulebrONT can help benchmark assembly pipelines, ONT genome sequencing datasets for four highly contrasted species were tested : *i*. a strain of *Acinetobacter baumannii* (Wick, Judd, et al., 2021) (ABJ9: reads N50 15,130 bp; median 7,621 bp; 741 Mb), *ii*. a strain of *Haemophilus haemolyticus* (Wick, Judd, et al., 2021) (HM1C1321: reads N50 10,569 bp; median 7,027 bp; 1,6 Gb) *iii. Meloidogyne graminicola* nematoda (Phan et al., 2020) (VN18: reads N50 9,372 bp; median 2,853 bp; 3.2 Gb) and *iv*. a *Oryza sativa* dataset (unpublished data) (DJ123: N50 22,328 bp, median 13 544 bp, total 14,6 Gb).

For bacterial sequences, all available assemblers included in CulebrONT were tested using reads for samples ABJ9 and HM1C1321 from *A. baumannii* and *H. haemolyticus*. For ABJ9, good results were obtained by Canu and Miniasm, but the *best* (the longest N50 and lowest L50) were found using Miniasm + 2 minipolish rounds + Medaka correction with only 2 circularised contig (N50 3,798,675 bp and L50 1) obtaining and a Busco score of 98.4%. Busco results obtained on this sample can be found on the figure 2. For HM1C1321, Flye gave the highest number of circularised contigs (N50 2,052,024 bp and L50 1) and the Busco score for conserved orthologous genes on the final assembly was 95.2%. More details can be found on the table 1).

**Table 1.**
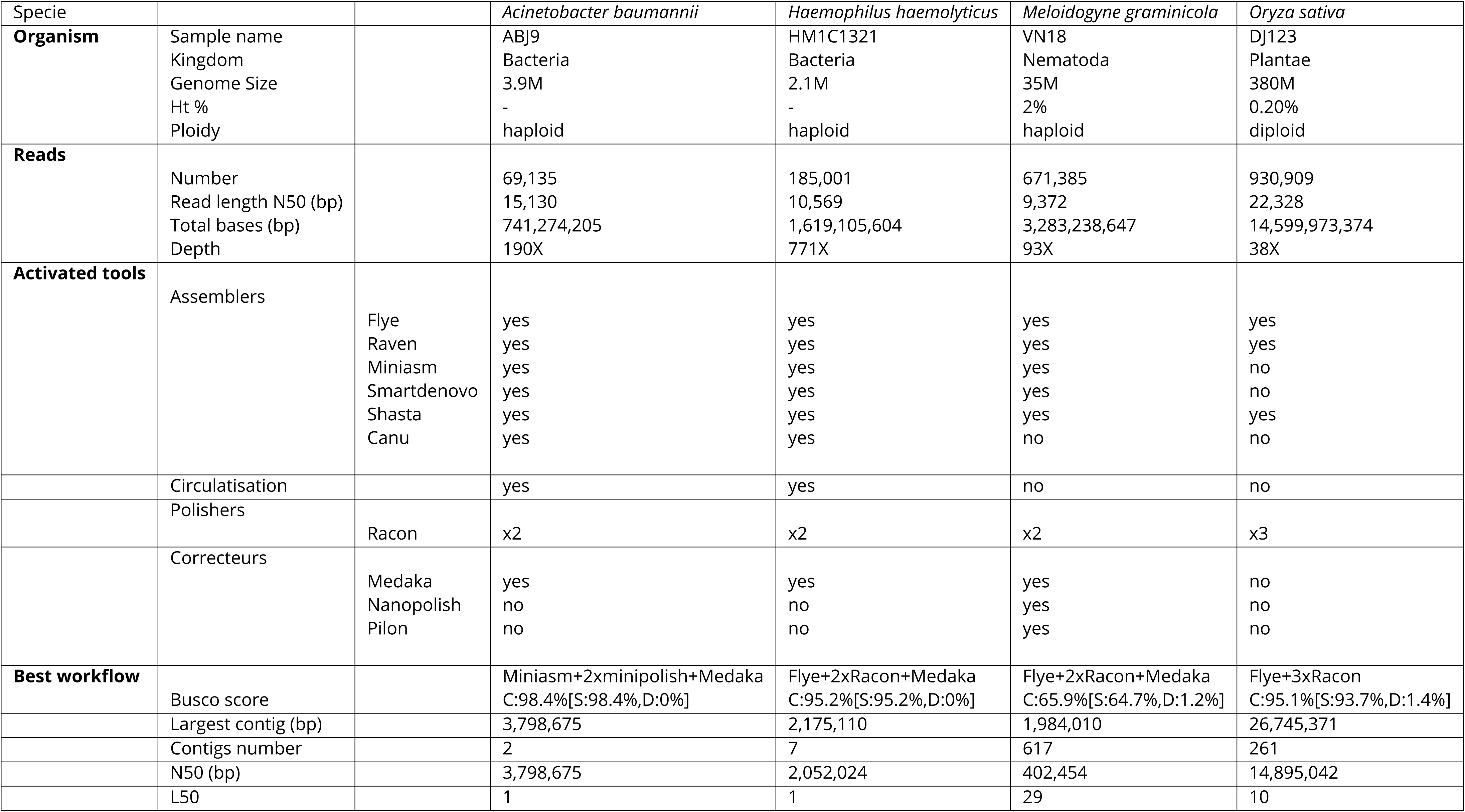
Results obtained on ONT data using CulebrONT on various biological models and assemblers. The *“Organisms”* row provides information about the samples, the *“Reads”* subsection describes input reads features, the *“Activated tools”* section lists the tools activated (‘yes’) in the corresponding CulebrONT run, and a summary of the best results is presented in the *“Best workflow”* row (considering best by the longest N50 and lowest L50 in this study).

Five assemblers were tested for *M. graminicola* (Flye, Shasta, Miniasm, Smartdenovo and Raven). With the VN18 sample, the lowest number of contigs (617) was obtained using a combination of Flye for assembly with a N50 of 402,454 bp and a L50 of 29 and followed by two rounds of Racon and ultimately Medaka polishing (see table 1).

On *Oryza sativa*, due to the larger genome size, only three assemblers were activated (Flye, Shasta and Raven) and polishing was performed with Racon only. For DJ123 rice sample, Flye + two Racon rounds seemed to be the optimal solution, with the best N50 (14,895,042 bp) and lower L50 (10 for 12 chromosomes). An overview of a CulebrONT report is shown on the figure 2.B using rice as an example. More details can be found on table 1 and more reports generated by CulebrONT are available at: https://itrop.ird.fr/culebront_utilities/PCI_RESULTS.

### Applications

CulebrONT has been also successfully used on many various organisms, including more than 40 bacteria (*Xanthomonas sp*) (S. Cunnac, pers com), haploid fungi (*Pseudocercospora fijiensis*) (J. Carlier, pers com) and (*Fusarium oxysporum f*.*sp. cubense*) (E. Wicker, pers com), green algae (*Bathycoccus sp*), insects (*Drosophila sp* (Mohamed et al., 2020)), diploid grasses (*Paspalum sp* (O. Blanc, pers com) and other *Oryza sp*), as well as on allopolyploid plants (ABB triploid banana; M. Rouard, pers com).

## Discussion

Obtaining a good assembly is a complex task, and the software ecosystem for this is evolving continually. While CulebrONT is currently proposing the state-of-the-art tools from today, we expect evolution in the next months or years for the next releases in different ways: On a short term, we will integrate the possibility of adding an external fasta sequence for using the QC part (for instance coming from an assembler not included in CulebrONT), re-entry at any steps, and also optimize the disk space usage.

On a medium term, we will re-evaluate the available tools (adding hifiAsm (Cheng et al., 2021), removing tools not updated for years), restructure the final output structure for more readability, and better integrate the outputs from diverse QC tools (in particular BlobTools or RAM usage per tool).

Finally, in a longer term, *i*.*e*. in the next 2 years, we plan to integrate tools for contig integrity, possibilities for haplotyping/polyploids and so on.

## Conclusion

In summary, CulebrONT simplifies the analysis of large-scale assemblies by allowing fast and reproducible processing of a single dataset or on a collection of samples simultaneously. The output facilitates the comparative evaluation of the assembly workflows while keeping a traceable record of the analyses.

## Acknowledgements

Data processing was performed at the High-Performance Computing Cluster i-Trop of IRD. Authors thank N.Tando for administration support and the South Green Plateform. We also thanks the French Bioinformatics Infrastructure IFB for feedback on CulebrONT installation.

Version 5 of this preprint has been peer-reviewed and recommended by Peer Community In Genomics (https://doi.org/10.24072/pci.genomics.100018)

## Data, script and code availability

CulebrONT pipeline project home page https://github.com/SouthGreenPlatform/CulebrONT_pipeline. Documentation : https://culebront-pipeline.readthedocs.io/en/latest/.

CulebrONT is also available as a Python Package https://pypi.org/project/culebrONT

Source code DOI : https://dataverse.ird.fr/dataset.xhtml?persistentId=doi:10.23708/TBPNWJ.

This pipeline is operating system platform independent.

CulebrONT has been programmed in Python >=3.6 langage and Snakemake >=5.10.

CulebrONT is under GPLv3 licence with any restrictions to use by non-academics.

## Author’s contributions

JO, FS and SC conceived the original idea. JO, AC, SR, FC, TV and SC written code source. Documentation was performed by JO, SR, FS, SC and AC. JO and FS analysed data. JO, FS and SC drafted the manuscript. All authors read and approved the final manuscript.

### List of abbreviations

N50: the smaller contig length required to cover 50 percent of the assembled genome sequence with the largest contigs.
L50: The number of sequences required to obtain N50.

## Supplementary information availability

Results generated by CulebrONT can be found in https://itrop.ird.fr/culebront_utilities/PCI_REPORTS/. Available data test can be found in https://itrop.ird.fr/culebront_utilities/Data-Xoo-sub/.

## Annexes or Supplementary Information

Additional file 1. Supplemental file descriptions of the CulebrONT implementation.

