## supplementary information for "CulebrONT: a streamlined long reads multi-assembler pipeline for prokaryotic and eukaryotic genomes"

July 13, 2022

#### 1 CulebrONT's architecture

CulebrONT is a snakemake pipeline originally created to assemble ONT reads, but also working on PB, that can also polish primary assemblies and provide quality metrics, in order to facilitate comparison of assembly methods. Using CulebrONT, workflows can be easily generated to assemble genome from various organisms, from bacteria to more complex eukaryots. Dedicated and optional steps are included to take into account the specific features of procaryotic genomes assembly, *i.e.*. circularization and definition of the origin of replication.

In order to build a pipeline, the user customizes mainly a file in YAML (Yet Another Markup Language) format: *config.yaml*. Two other files may be necessary to install dependencies *tools_path.yaml* and manage resources on a HPC with a job scheduler environment *cluster_config.yaml*. CulebrONT uses the snakemake architecture in order to run a relevant set of rules, each one being executed into a dedicated virtual environment.

##### 1.1 *config.yaml*

The main workflow parameters determining the assembly strategy are stored and read by CulebrONT from *config.yaml* (see figure 1.A). This file provides the path to the fastq (required, plain or gzipped), fast5 and reference genome (if any) files. For each step (assembly, polishing or correction), the user can choose which tools, including quality evaluation tools, must be launched and can define custom options (or default) for these programs. This allows users to run different workflows on exactly the same dataset, while recording tools versions and parameters.

##### 1.2 *tools_path.yaml*

CulebrONT uses singularity to setup dependencies of each workflow (see figure 1.B). Pre built images are available on the i-Trop IRD platform server `https://itrop.ird.fr/culebront_utilities/singularity_build/`. It is also possible to build singularity images on your local or HPC machines, but root access is necessary. Dedicated singularity recipes can be found on our github repository (`https://github.com/SouthGreenPlatform/CulebrONT_pipeline`). These *Singularity recipes* containers allows an easy installation of CulebrONT dependencies tools. Alternatively, CulebrONT also allows the use of Environment Modules usually found on clusters.

The *tools_path.yaml* file specifies the paths of the singularity containers or of the available module environments.

##### 1.3 *cluster_config.yaml*

Within CulebrONT, users can take full advantage of Snakemake for workflow execution on a cluster or on the cloud.

The *cluster_config.yaml* file defines the requested HPC resources (memory, CPU number, partitions, CPU/GPU access) for executing rules on a cluster with a job scheduler, and can be adapted by the user or system administrators (see figure 1.C). If needed, execution through command-line cluster wrappers provide additional controls and customization possibilities to the user. An easy-to-use Snakemake profile method is described on our documentation page. In the absence of a cluster setup, the workflow will be executed locally.

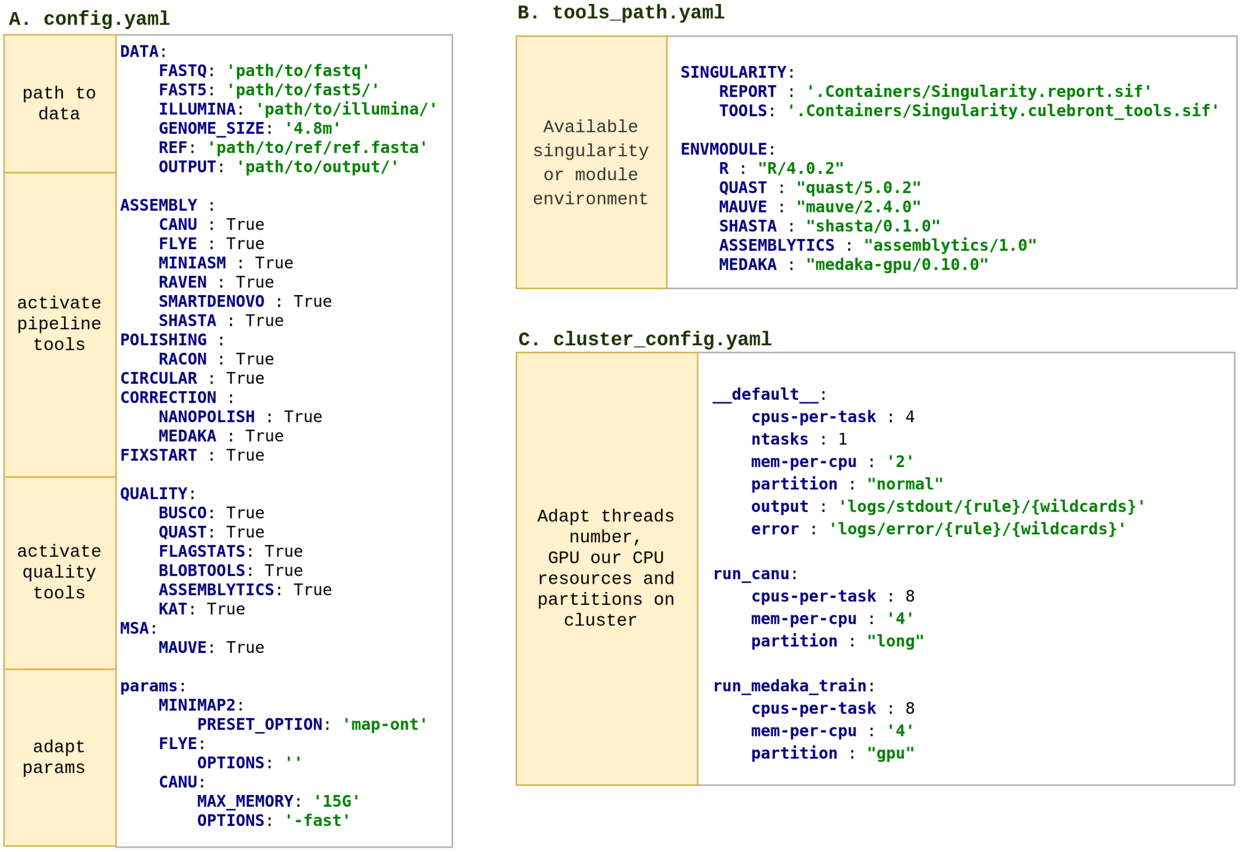


Figure 1: Configuration files setup defining an instance of the workflow.

### 2 How to run CulebrONT?

A comprehensive documentation can be found online `https://culebront-pipeline.readthedocs.io/en/latest/` and a test dataset is available on the i-Trop server `https://itrop.ird.fr/culebront_utilities/`

#### 2.1 Building pipelines from *config.yaml*

We present here some examples of modular pipelines build using the *config.yaml* file (figure 2). Users can be interested to test the best assembly pipeline on a bacterial reference strain before launching it on multiple related strains, for instance. If an eukaryotic organism with a large genome size ($\geq$ 100Mb or more) is targeted, starting with testing all assemblers and checking quality of raw assemblies before relaunching CulebrONT with polishing and correction with only the best assembler(s) is more relevant. The user can thus choose optional tools for each integrated step.

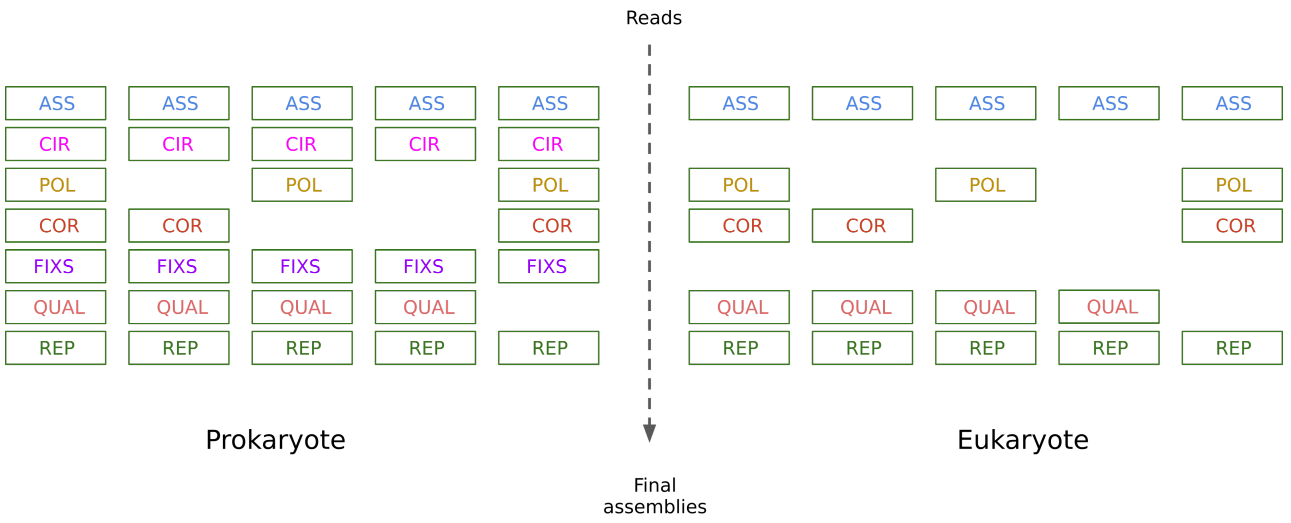


Figure 2: Several possibilities to build a workflow. ASS: assembly, CIR: circularisation, POL: polishing, CORR: correction, FIXS: fixstart, QUAL: quality control, REP: final report. For each step, multiple tools are available.

#### 2.2 Building a pipeline for prokaryotes or eukaryotes

**Assembly**: CulebrONT currently includes six recent and community-validated assemblers (Figures 4 and 3, blue squares): Flye[1], Canu[2], Miniasm[3] + Minipolish[4], Shasta [5], Smartdenovo[6] and Raven[7]. Tools versions for each release can be found in the CulebrONT documentation.

- Canu[1] performs assembly by using corrected and trimmed reads. Its assembly strategy uses a overlap-layout-consensus (OLC) approach.
- Flye[2] generates the concatenation of multiple disjoint genomic segments called disjointigs to build a repeat graph. Reads are mapped to this repeat graph to resolve conflicts (unbridged repeats) and output contigs.
- Miniasm[3] performs the layout step of OLC. Read overlapping is performed separately with Minimap2[8].
- Raven[5] also uses an OLC approach. The overlapping step is similar to Miniasm and performed by Minimap2. The initial consensus step uses Racon. Layout step removes spurious overlaps from the graph, improving contiguity.
- Smartdenovo[6] performs read overlapping, rescues missing overlaps, identifies low-quality regions and produces a consensus.
- Shasta[7] is a computationally efficient assembler, relying on a reduced representation of marker k-mers used to find overlaps and to build an assembly graph.

[1] `https://github.com/fenderglass/Flye`
[2] `https://canu.readthedocs.io/en/latest/`
[3] `https://github.com/lh3/miniasm`
[4] `https://github.com/rrwick/Minipolish`
[5] `https://github.com/chanzuckerberg/shasta`
[6] `https://github.com/ruanjue/smartdenovo`
[7] `https://github.com/lbcb-sci/raven`
[8] `https://github.com/lh3/minimap2`

**Polishing**: This step is ensured by Racon[9][8] (Figures 4 and 3, yellow squares). Racon corrects raw contigs generated by raw assembly methods using the original ONT or PB reads. The user can specify a number of Racon rounds (constrains from 1 to 9 rounds) and CulebrONT will recursively execute them (generally between 2 to 4 iterations are commonly done). Polishing from assembled pseudomolecules obtained using Miniasm is automatically performed by Racon, also included in minipolish.

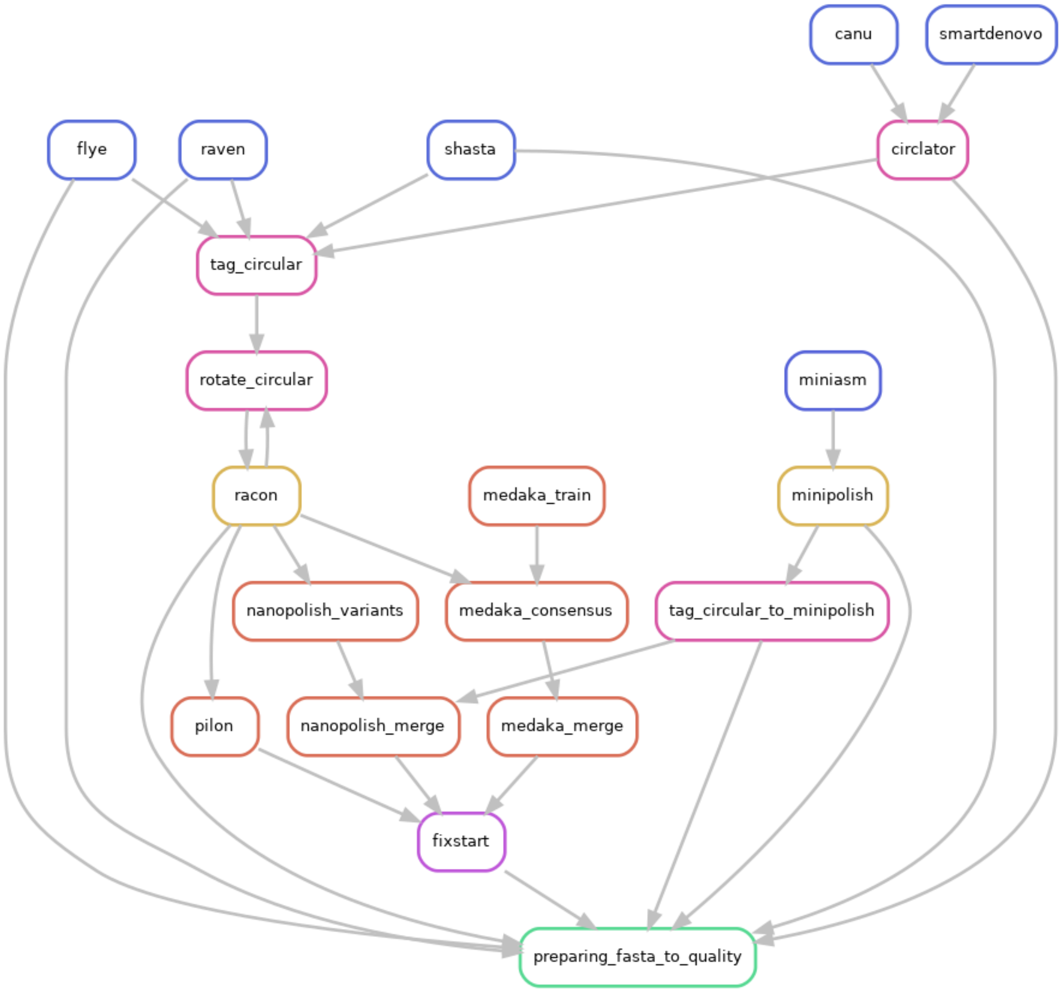


Figure 3: Circular option on *config.yaml* file is activated

**Correction**: Correction can improve the consensus sequence for a draft or polished ONT-based genome assembly. Community validated tools Nanopolish[10] and Medaka[11] are included (Figures 3 and 4, orange squares). Medaka uses neural networks applied to a pileup of individual sequencing reads in fastq against a draft assembly. The model training can be performed using CulebrONT or provided externally. Correction by segments of the assembled molecules is manage by CulebrONT, for both Nanopolish and Medaka, in a parallel way, improving speed.

Pilon[12] has been recently included and can be activated in order to correct assemblies using Illumina short reads.

[9] `https://github.com/isovic/racon`
[10] `https://nanopolish.readthedocs.io/en/latest/index.html`
[11] `https://github.com/nanoporetech/medaka`
[12] `https://github.com/broadinstitute/pilon/wiki`

Even if prokaryotic genomes are smaller and less repetitive than eukaryotic ones, **Circularisation** represents a specific dimension of the assembly task. If an assembled molecule, for example a bacterial chromosome or a plasmid, is identified as circular, this molecule is tagged and will be specially treated in the pipeline (see figure 3, pink squares). If the circular step is activated, the $--plasmids$ option on Flye (<=v2.9) will be automatically used. For Canu and Smartdenovo, Circlator[13] is used to circularise assemblies. Circularisation for Miniasm is performed by Minipolish[14][4], which first polishes Miniasm graph assembly using Racon and then handles circularisation. In CulebrONT, for others assemblers (not miniasm), tagging and rotation of circular molecule before each polishing step is implemented. The starting position of circular contigs is arbitrarily shifted between polishing rounds. The circular status of Miniasm, Raven and Shasta contigs is determined by inspecting assemblers GFA files, or a log file for Flye and Circlator, in a dedicated *tag_circular* step, where only circular fasta sequences are tagged.

The **Fixstart** rule uses the fasta file generated after the most downstream step of the pipeline. It shifts the start position of circular sequences so that the origin of replication of the contig coincides with the start of the ortholog of the *dnaA* gene (if found). This is useful to harmonize the origin of bacterial genomes in a multiple alignment.

The circular option can be *True* or *False* in the *config.yaml* file. The Figure 4 presents a possible pipeline without the circular step, for eukaryotes for instance.

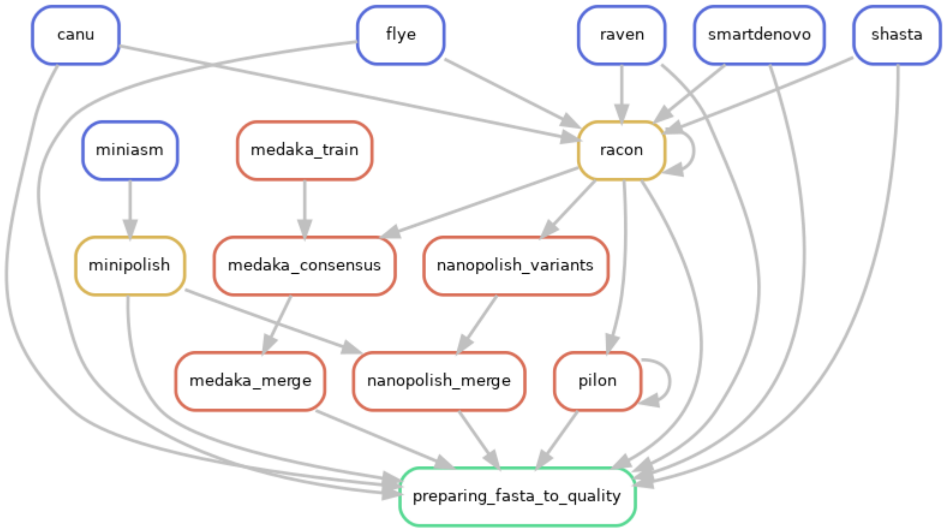


Figure 4: Circular option on *config.yaml* file is deactivated

#### 2.3 Quality control

A wide variety of quality control tools is implemented in CulebrONT to check accuracy of a given assembly (all optional). *Busco*[9] evaluates genome completeness and accuracy by comparison of predicted genes with a specified database of conserved genes. Various metrics, such as the number of contigs, length of the longest contig, N50, number of predicted genes, numbers of potential misassemblies are computed using *Quast*[10]. If *Busco* or *Quast* are selected, their statistics will be computed for all assemblies generated by the pipeline, regardless of their stage in the workflow.

[13] `https://github.com/sanger-pathogens/circlator`
[14] `https://github.com/rrwick/Minipolish`

Their statistics are summarized in the CulebrONT final HTML report. The complete output of these tools can also be explored in the output directory.

Supplemental tools are also included: *blobtools*[11] to detect contamination, *assemblytics*[12] to compare contiguity of the assemblies against a reference genome, *KAT*[15] and *Merqury*[16] to explore k-mers frequencies and check possible contamination, samtools FLAGSTATS to compute remapping percentage over assemblies, and *Mauve*[13] to generate a multiple-alignment of the alternative assemblies (for small bacterial genomes). Only the fasta file generated after the most downstream step of the pipeline is analyzed by these optional tools. If selected, their statistics are also summarized in the CulebrONT final HTML report.

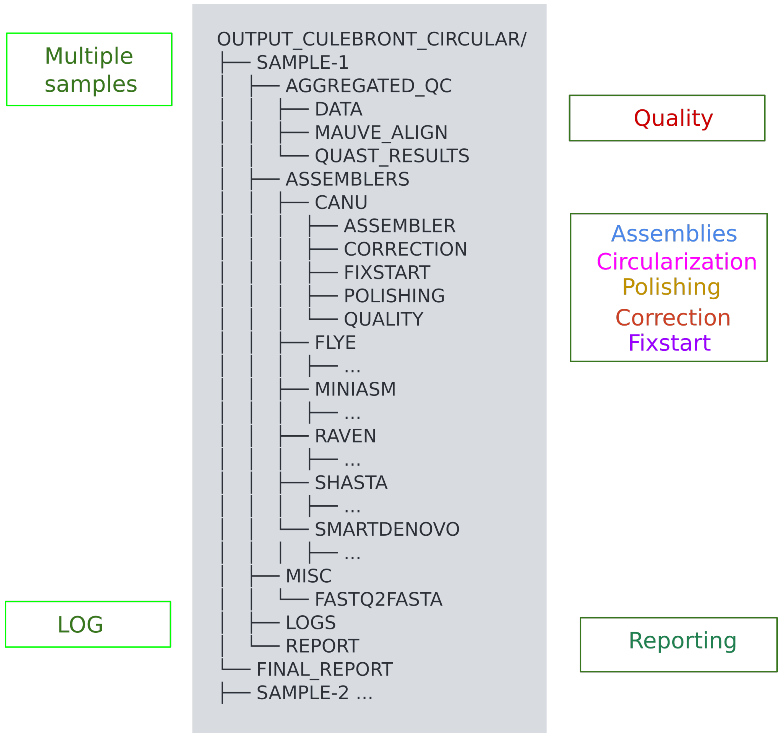


Figure 5: Output folder architecture

#### 2.4 Output and reporting

CulebrONT generates an output directory with a specific architecture (Figure 5). For each analysed sample, the output directory contains subdirectories named after selected steps in the *config.yaml* file (assembly, circularization, polishing, correction, fixstart, and quality control). In addition, a **LOG** directory contains all logs created. In the **FINAL_REPORT** folder, CulebrONT generates an HTML report, summarizing statistics from the various steps of the pipeline and including a record of the configuration parameters used in the pipeline, the version of the software tools and the statistics about computing time and resources. An example of the report generated by the test dataset can be found here: `https://itrop.ird.fr/culebront_utilities/FINAL_REPORT/CulebrONT_report.html`.

[15] `https://github.com/TGAC/KAT`
[16] `https://github.com/marbl/merqury`

##### 2.5 Installation using a PyPi package

CulebrONT is available as a Pypi package `https://pypi.org/project/culebrONT/`. This Python API allows users easy install CulebrONT in local or HPC machines `https://culebront-pipeline.readthedocs.io/en/latest/`.

##### 2.6 Conclusion

CulebrONT can be used in many experiments and on many samples, and in addition is a FAIR product :

- Open source: You can find it on github with a licence under CeCill-C (`http://www.cecill.info/licences/Licence_CeCILL-C_V1-en.html`) and GPLv3. You can modify and redistribute it.
- Modulable: All steps to obtain the best quality assembly are included, and can be activated (or not) according to your needs. All the most commonly used tools in the community are integrated, as well as various state-of-the-art quality control tools. The possibility to use containers makes it easy to change and control tools version.
- Scalable: most eukaryote or prokaryote genome can be assembled and circular molecules are handled accordingly. You just need to fill in a simple text-based configuration file to build a personalised pipeline. The limits are imposed by the hardware, not the code.
- Intuitive: in order to quickly give the user an idea of the best final assembly, a well structured output directory and log files are provided as well as a report with statistics and images of the most important steps.
- Traceable: users can reproduce the analysis easily using the report and config.yaml, and the HTML report will provide all information about softwares version, original data, QC, and options used.

##### 2.7 Related pipeline of interest

BaseDMux[17] is another snakemake pipeline developed in collaboration with the CulebrONT projet. It allows basecalling, demultiplexing and filtering ONT data in a flexible way before assemblies. Its output can be used directly as input for CulebrONT.

##### 2.8 Competing interests

The authors declare no competing interests.

[17] `https://github.com/vibaotram/baseDmux`

[4] RR. Wick and KE. Holt. Benchmarking of long-read assemblers for prokaryote whole genome sequencing [version 3; peer review: 4 approved]. *F1000Research*, 8(2138), 2020.

[5] R. Vaser and M. Šikić. Raven: a de novo genome assembler for long reads. *bioRxiv*, 2020.

[6] https://github.com/ruanjue/smartdenovo
